# An empirical test of the role of value certainty in decision making

**DOI:** 10.1101/2020.06.16.155234

**Authors:** Douglas Lee, Giorgio Coricelli

**Affiliations:** University of Southern California, Department of Economics

## Abstract

Most contemporary models of value-based decisions are built on value estimates that are typically self-reported by the decision maker. Such models have been successful in accounting for choice accuracy and response time, and more recently choice confidence. The fundamental driver of such models is choice difficulty, which is almost always defined as the absolute value difference between the subjective value ratings of the options in a choice set. Yet a decision maker is not necessarily able to provide a value estimate with the same degree of certainty for each option that he encounters. We propose that choice difficulty is determined not only by absolute value distance of choice options, but also by their value certainty. In this study, we first demonstrate the reliability of the concept of an option-specific value certainty using three different experimental measures. We then demonstrate the influence that value certainty has on choice, including accuracy (consistency), choice confidence, response time, and choice-induced preference change (i.e., the degree to which value estimates change from pre- to post-choice evaluation). We conclude with a suggestion of how popular contemporary models of choice (e.g., race model, drift-diffusion model) could be improved by including option-specific value certainty as one of their inputs.

## INTRODUCTION

For many decades, researchers in economics, psychology, and cognitive neuroscience have studied the concept of subjective value, and how implicit subjective value influences explicit choices. In more recent years, decision researchers have frequently relied on self-reported estimates of subjective value (value ratings) to support their theories and models (see Rangel et al, 2008, for a review). Value ratings collected by self-report have served as the fundamental input of most contemporary models of choice. One key component of many models is what is referred to as “choice difficulty”, which is most often defined as the distance between the subjective value ratings of the options in the choice set (where difficulty declines with distance). Choice difficulty has been shown to reliably predict both choice (i.e., the probability of choosing the higher-rated option decreases with difficulty) and reaction time (i.e., deliberation time increases with difficulty) (e.g., Hunt et al, 2012; Palmer et al, 2005). Yet until recently, most models did not explicitly incorporate the possibility that a decision maker (DM) might not be fully certain about the value ratings that he reports. For example, a DM might have an estimate about the (subjective) value of an option, yet simultaneously have a belief about the accuracy of his value estimate. Said another way, sometimes we might feel that we like something a certain amount, but also feel more or less sure about that amount—sometimes we might feel that we know for sure precisely how much we value something, other times we might feel unsure about our value estimates. Consequently, when choosing between items, certainty about the values of the individual options directly impacts choice confidence, which can be defined as a feeling of certainty about which of the options has a higher value. The more uncertain the individual option values are, the more difficult it is to determine which one has the higher value. It would therefore follow that a more complete definition of choice difficulty should account for such a concept as subjective certainty.

Some recent studies have indeed included value certainty as an independent variable in their models. DeMartino et al (2013) examined the effects of what they called “bid confidence” on the dependent variables in their model. The authors suggested that bid certainty should moderate the impact of choice difficulty (defined as value rating distance) on choice accuracy, choice confidence, and choice response time (RT). Lee and Daunizeau (2020a) took this a step further, demonstrating in their model how value certainty might be explicitly involved in choice. In that study, the authors suggested that when value ratings are uncertain, a DM will be less accurate and less confident in his choice, but he will also be more inclined to invest mental effort (for which RT could serve as a proxy) in order to decrease the value uncertainty and enable him to confidently choose his preferred option. In fact, in that model, one of the principal drivers of the proposed effort-confidence tradeoff is the desire on the part of the DM to increase the certainty that he has about his value estimates. The model builds upon previous work that demonstrated the same principle in a less formal manner, by showing that both RT and preference change (difference between post- and pre-choice ratings) were decreasing functions of value certainty (for which the authors used the average value certainty of the options being compared), while choice confidence was an increasing function of value certainty (Lee and Daunizeau, 2020a). In their model, the authors explain that lower value certainty impairs the ability of the DM to distinguish the options. This dampens choice confidence, which the DM attempts to boost through mental effort (proxied by RT). In turn, the effort allocation leads to value estimate refinements, and potentially changes of mind (preference reversals).

Recent work in other, non-subjective value-based domains also suggests that a measure of certainty about the options is conceptually important when examining choice behavior (Pouget et al, 2016). Frydman and Jin (2019) invoke the principle of efficient coding to suggest a link between value certainty and choice behavior. In this study, certainty spawns from repeated exposure, which causes greater precision in neural representation (i.e., efficient coding). Higher certainty, thus defined, leads to higher choices accuracy (i.e., consistency with value estimates). Padoa-Schioppa and Rustichini (2014) illustrate a similar concept based on *adaptive* coding, where neural activity is normalized according to the range of option values in the current environment, thus causing choice stochasticity to increase as representation precision decreases. Along similar lines, Woodford (2019) explains how many important aspects of economic choice (e.g., choice stochasticity, risk aversion, decoy effects) could result from noisy neural representations of value. Although the author does not directly refer to a subjective feeling of certainty, it is no far stretch to relate the precision of neural representations to a subjective feeling of certainty. In Polania et al (2019), the authors refer to efficient coding as well as Bayesian decoding principles to explain how choice behavior is influenced by value certainty. Interestingly, the authors do not record any self-reports of value certainty to validate their model. Instead, certainty in this study is captured by rating consistency (i.e., similarity of repeatedly self-reported value estimates). The basic idea here is that greater precision in the neural encoding of value will lead to greater consistency across multiple interrogations of the value-encoding neural population. This precision thus leads to both more consistent ratings and more choices consistent with those ratings.

In spite of recent theoretical and empirical evidence that value certainty plays an important role in choice behavior, most popular models of value-based decision making still do not include this variable. In particular, so-called accumulation-to-bound models such as the race model (e.g., Tajima et al, 2019; Kepecs et al, 2008) and the drift-diffusion model (e.g., Kiani and Shadlen, 2009; Krajbich and Rangel, 2011) do not include option-specific certainty. Tajima et al (2019) model noise in the accumulation process at the system level, where all options have the same degree of uncertainty imposed upon them by the environment, rather than at the option level (although the authors themselves suggest that future studies should explore the various sources of value uncertainty). Other studies have similarly included systemic, but not option-specific, uncertainty (e.g., Louie et al, 2013). Krajbich and Rangel (2011), along with most other published versions of the drift-diffusion model (DDM), fail to include a variable to represent value (un)certainty.

Perhaps more researchers would be willing to include value certainty in their models if there was more available evidence demonstrating that certainty could be reliably measured. In this study, we hope to provide some such evidence. In line with Lee and Daunizeau (2020a, 2020b), we explicitly ask decision makers to report their subjective feelings of certainty about the ratings they provide about each of a large set of options. In line with Polania et al (2019), we also implicitly capture value certainty by calculating consistency across multiple ratings of the same options. We then show that the explicit and implicit measurements of value certainty are highly correlated for each individual DM, which suggests that they are both reliably expressing the same internal representation precision. We show that RT during value estimation for each item is also strongly correlated with both measures of certainty, which suggests a link between the representation precision and the cognitive effort required to decode the value signal. Finally, we show that self-reported estimates of certainty generally increase across repeated value estimations. This should be expected if contemplating the value of an option is tantamount to constructing its internal representation, because as the cumulative total of processed information rises (i.e., across multiple rating sessions), the precision of the representation should also increase.

In a second study, which is essentially a replication of the rating-choice-rating paradigm of Lee and Daunizeau (2020a, 2020b), we reproduce the previous findings that value estimate certainty positively influences choice consistency and choice confidence, and negatively influences response time and choice-induced preference change. We also show an interesting novel result—we can predict which option a DM will choose based on the relative value estimate certainty of the options being compared, even while ignoring the options’ value estimates themselves.

In sum, we suggest that researchers should no longer neglect the concept of value certainty when building their models. We provide confirmatory and novel evidence that value certainty plays an important role in the cognitive process of decision making. In particular, we suggest that of the three different measures of value estimate certainty that we examined (self-reports, rating consistency, rating RT), self-reported ratings are ideal, but that consistency could work well if multiple ratings are available for each item. With respect to using rating RT as a proxy for value certainty, we suggest that this could suffice if no other measures were available, but that caution should be exercised when interpreting the results.

Note: for clarity, we explicitly use different terminology throughout this paper for subjective beliefs about value estimates and about choices: “certainty” refers to the subjective feeling of certainty about a value estimate rating; “confidence” refers to the subjective feeling of confidence that the chosen item is the better one. We never use these terms interchangeably.

## METHODS

We conducted a pair of behavioral experiments with the intention of demonstrating the reliability of various measures of value certainty, which include: self-reports, rating consistency, and response time. Participants considered a set of 200 options and provided three separate value ratings for each option, as well as three separate certainty reports about those ratings. We also recorded response time for each evaluation. Participants also made choices between pairs of options, as well as choice confidence reports. Detailed task descriptions can be found below.

In Study 1, we asked participants to rate the value of each of a series of items. In addition to the standard subjective value question, we also asked participants to rate their subjective certainty regarding each subjective value judgment. Because we are interested in assessing the consistency of value ratings and how that relates to subjective certainty ratings, we repeated the value and certainty ratings three times during the experiment. This allows us to assess each of our hypotheses, in particular: the correlation between value rating consistency and certainty rating; the increase in certainty across repeated value ratings; the decrease in response time across repeated value ratings.

In Study 2, we asked a different group of participants to rate the value of each of a series of items, as well as to make choices between pairs of items. In addition to the standard subjective value question, we also asked participants to rate their subjective certainty regarding each subjective value judgment, and their subjective confidence regarding each choice.

### Materials

We built our experiment in Gorilla (gorilla.sc). The experimental stimuli consisted of 200 digital images, each representing a distinct snack item food item. The stimulus set included a wide variety of items. Prior to commencing the experiment, participants received a written description about the tasks and detailed instructions on how to perform them.

### Ethics statement

Our analysis involved de-identified participant data and was approved by the ethics committee of the University of Southern California (USC). In accordance with the Helsinki declaration, all participants gave informed consent prior to commencing the experiment.

### Study 1

#### Participants

A total of 37 people participated in this study (22 female; age: mean=21.4, stdev=3.6, min=18, max=35). All participants were recruited from the undergraduate population at USC using SonaSystems. Each participant received course credit as compensation for one hour of time.

#### Experimental Design

The experiment was divided into four sections. There was no time limit for the overall experiment, nor for the different sections, nor for the individual trials.

In the first section (Exposure), participants merely observed as all individual items were displayed in a random sequence for 750ms each. The purpose of the Exposure section was to familiarize the participants with the full set of items that they would later evaluate, allowing them to form an impression of the range of subjective value for the set. This would diminish the possibility that value ratings would become more accurate across time merely due to a dynamic adaptation of the range as the participants viewed new items.

The second through fourth sections (Rating1, Rating2, Rating3) were identical, except for the sequence of trials. In each rating section, all stimuli were displayed on the screen, one at a time, in a random sequence (randomized across participants and across sections for each participant). At the onset of each trial, a fixation cross appeared at the center of the screen for 750ms. Next, a solitary image of a food item appeared at the center of the screen. Participants responded to the question, “How pleased would you be to eat this?” using a horizontal slider scale. The leftmost end of the scale was labeled “Not at all.” and the rightmost end was labeled “Very much!” The scale appeared to participants to be continuous, and the data was captured in increments of 1 (ranging from 1 to 100). Participants could revise their rating as many times as they liked before finalizing it. Participants clicked the “Enter” button to finalize their value rating response and proceed to the next screen. Participants then responded to the question, “How sure are you about that?” by clicking on a horizontal Likert scale (left to right: “not at all”, “slightly”, “somewhat”, “fairly”, “very”, “extremely”) to indicate their level of subjective certainty regarding the preceding value judgment. At that time, the next trial began. Participants were not aware that there were three different rating sections, as the design technically only included one rating section. Within that section, the sequence of trials included a random ordering of all items, followed by another random ordering of all items, followed by another random ordering of all items. From the perspective of the participant, the study consisted of a long series (600) of item evaluations, where repetition could (and indeed, always did) occur. This design, in which participants could not anticipate that each item was going to be rated multiple times, helped to preclude participants from explicitly trying to remember and replicate their evaluations across repetitions. A typical within-trial event sequence is shown in Figure 1.

**Figure 1:**
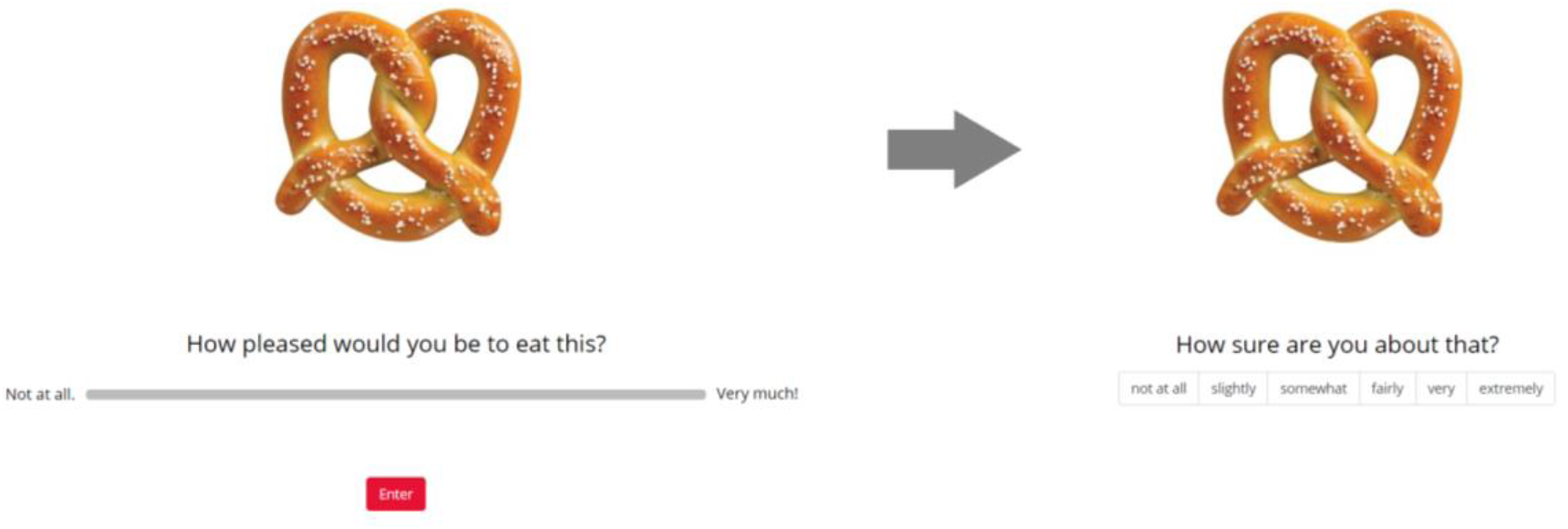
an example of a within-trial even sequence for the rating tasks (study 1 and study 2)

### Study 2

#### Participants

A total of 50 people participated in this study (18 female; age: mean=30.5, stdev=11.4, min=18, max=64; 8 missing gender info; 2 missing age info). All participants were recruited from the online subject pool using Prolific (prolific.co). Each participant received $5 as compensation for 45 minutes of time.

#### Experimental Design

The experiment contained three different tasks: exposure, rating, and choice. There was no time limit for the overall experiment, nor for the different tasks, nor for the individual trials.

The Exposure task was structurally identical to that described above for Study 1.

The Rating1 and Rating2 tasks were identical, except for the sequence of trials. For each rating task, all of the stimuli to which the participant had initially been exposed were again displayed on the screen, one at a time, in a random sequence (randomized across participants and across sections for each participant). The structure of the rating trials was identical to that described above for Study 1. Prior to commencing Rating2, participants were reminded that they should report what they felt at that time and not try to remember what they reported during Rating1. This helped to preclude participants from explicitly trying to remember and replicate their evaluations across repetitions. A typical within-trial event sequence is shown in Figure 1 above.

For the Choice task, all stimuli were displayed on the screen, two at a time, in a random sequence (randomized across participants and across sections for each participant). At the onset of each trial, a fixation cross appeared at the center of the screen for 750ms. Next, a pair of images of food items appeared on the screen, one towards the left, one towards the right. Participants responded to the question, “Which would you prefer to eat?” by clicking on the image of their preferred item. Participants then responded to the question, “Are you sure about your choice?” using a horizontal slider scale. The leftmost end of the scale was labeled “Not at all!” and the rightmost end was labeled “Absolutely!” Participants could revise their confidence report as many times as they liked before finalizing it. Participants clicked the “Enter” button to finalize their confidence report and proceed to the next screen. A typical within-trial event sequence is shown in Figure 2.

**Figure 2:**
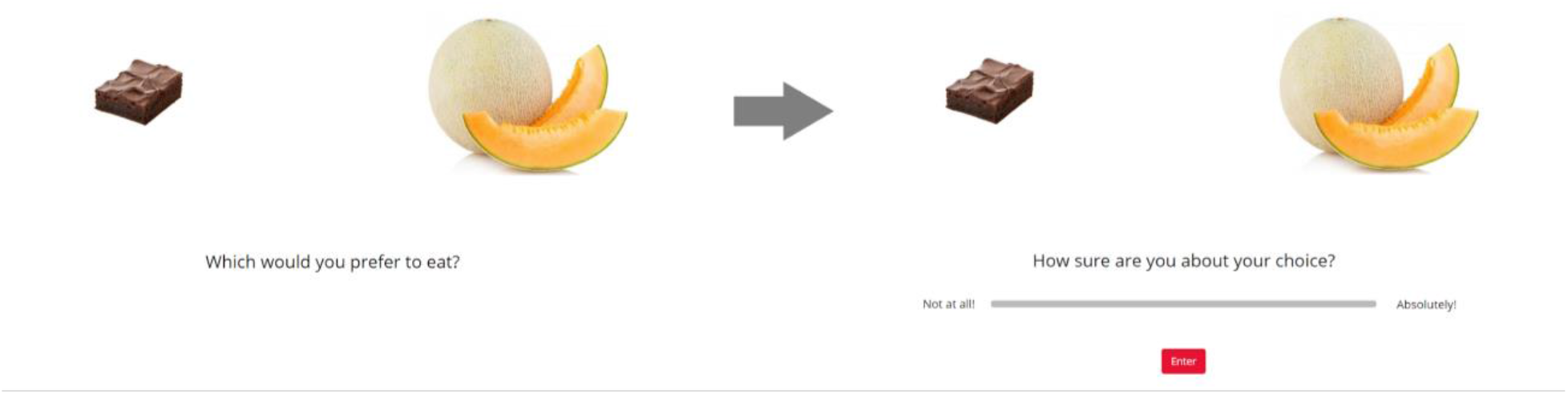
an example of a within-trial even sequence for the choice task (study 2)

The pairings of items for each choice trial were created in a deliberate manner. Specifically, we wanted to maximize the number of difficult choices that participants would be faced with. Here we define difficulty as the similarity of the value ratings between choice pair items. Because our simplified online experimental design did not allow for choice pairs to be created dynamically based on each participant’s personal subjective value ratings, we relied on our data from Study 1. That data provided us with value ratings for 200 items across 37 participants, which we used to calculate population statistics (median and variance of value estimate ratings) for each item. We first calculated the population value estimate variability, which was the variance of the value estimate ratings for each item across all 37 participants. Because we only wanted 150 items for Study 2, we sorted the original 200 items from lowest to highest population variance and removed the 50 highest-variability items (i.e., the items for which different participants had provided the most variable value estimate ratings) from our set. We thought that this would improve our chances that a new participant would rate the items similarly to the population average ratings. Next, we calculated the population median value for each item. We used the median instead of the mean so as not to be unduly influenced by extreme ratings. Sorting the item set from highest to lowest value, we created triplets of items (i.e., [item1 item2 item3], [item4 item5 item6], …). We created 50 choice pairs for by selecting the first and second elements from each triplet. We created an additional 50 choice pairs by selecting the first and third elements from each triplet. We thus had a total of 100 choice pairs, all of which should be difficult trials based on population statistics. (The reason why we created two separate sets in the manner described was to allow us to pilot test a hypothesis for a future study, but is irrelevant to this current study.) Obviously, individual ratings deviate from population ratings, which would naturally cause many of the choice pair trials to be more or less difficult for individual participants.

The usage of population median ratings in this way was solely to create choice pairs that would *a priori* be likely to be difficult for most participants—it had no impact on the choice data itself, which was based on the individual value estimate ratings of the participants who would actually make the choices. Therefore, although choice pairs were created based on population value estimate ratings from Study 1, the actual choice difficulty analyzed in the data for Study 2 was determined entirely by the personal value ratings provided by each participant in that study. Fortunately, this technique did indeed result in each participant in Study 2 facing a large number of difficult choices (defined by their own personal value ratings). Note: the validity of our analysis would not have been impacted either way, but the effects of interest would likely have diminished.

## RESULTS

Before conducting our main analyses, we first validated that our data were reliable (see Supplementary Material for details). We determined that the data were generally reliable, although we decided to exclude 11 participants from Study 1 and six participants from Study 2, for failing to perform the tasks properly for the duration of the experiment (see Supplementary Material for details).

### Study 1

#### Hypothesis 1: Certainty should negatively correlate with rating inconsistency

Certainty reports were provided by the participants during the study, but we needed to obtain a measure of rating inconsistency. For each participant, we thus calculated the within-item across-section variance of value ratings (i.e., V[Rating1_i Rating2_i Rating3_i] for i=1:200). We deemed that variance is a measure of inconsistency, because perfect consistency would yield a variance of zero and higher degrees of inconsistency would yield higher variance. For each participant, we used the average certainty for each item across the three rating sections as our measure of certainty. The correlation between certainty and inconsistency was negative and significant, as expected (median Spearman’s rho = −0.245, p<0.001). (See Figure 3.)

**Figure 3:**
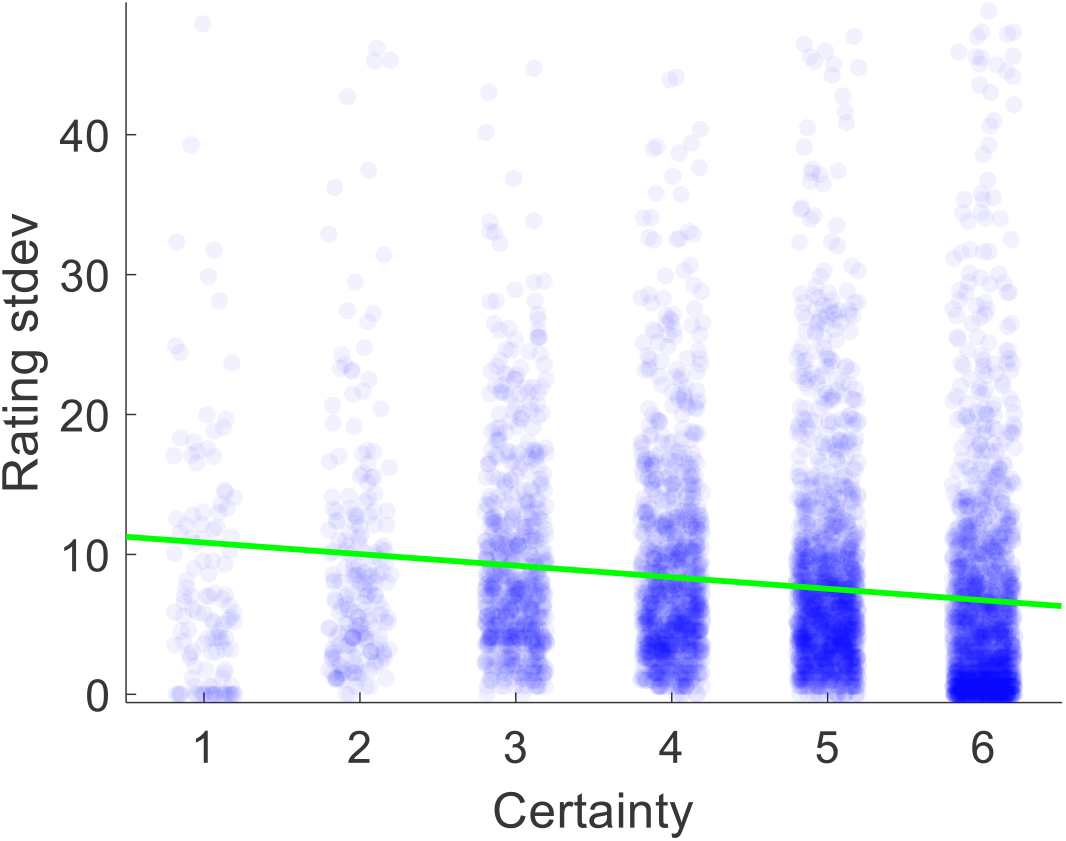
Scatter plot of the relationship between value estimate certainty and the standard deviation of ratings across rounds 1-3, pooled across all participants. Each dot represents one trial.

#### Hypothesis 2: Certainty should increase with repeated ratings

We first calculated the within-participant mean of certainty reports separately for Rating1, Rating2, and Rating3. We then calculated the group averages for these values. The across-participant across-item mean certainty for Rating1, Rating2, and Rating3 was 4.67, 4.74, and 4.84, respectively (see Figure 4). The increase in average certainty between Rating1 and Rating2 and between Rating1 and Rating3 were marginally significant (p=0.086, p=0.068; two-sided t-tests), but the increase in certainty between Rating2 and Rating3 was not significant (p=0.198; two-sided t-test).

**Figure 4:**
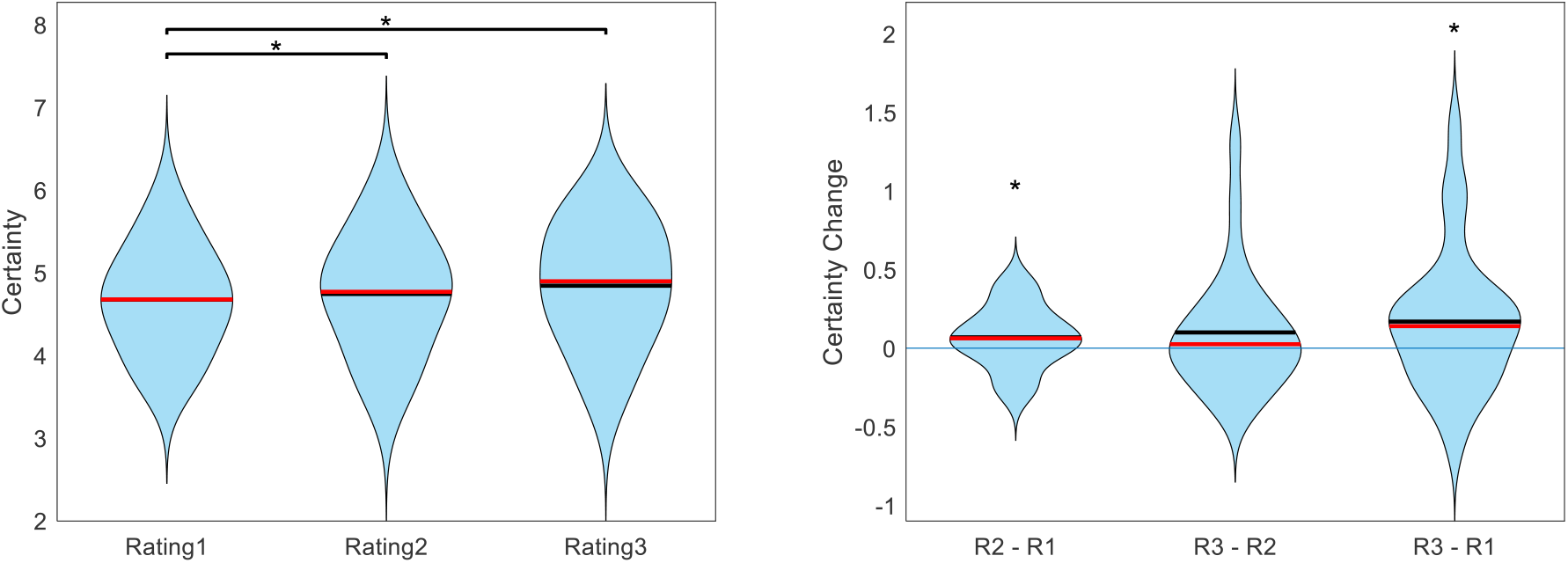
Across participants, value estimate certainty (across items) increased with each additional rating. Figure shows the cross-participant mean of within-participant mean certainty ratings across trials, separate for each rating task. (black lines indicate means, red lines indicate medians, significance stars indicate: * p<0.05, ** p<0.01, *** p<0.001)

In addition to the gradual increase in average certainty from Rating1 to Rating3, we also checked to see if there was a gradual decrease in average response time (RT). Because online testing is often plagued by distractions that cause some trials to have exceptionally long response times, we first removed all outlier trials. We defined an outlier as any trial in which RT was greater than the within-participant median RT plus three times the within-participant median average deviation. This resulted in an average of 9 trials being removed per subject (out of 100). After cleaning the data in this way, we indeed found that average RT decreased from one rating section to the next. The across-participant across-item mean RT for Rating1, Rating2, and Rating3 was 4.75s, 3.77s, and 3.58s, respectively (see Figure 5). The decrease in RT between Rating1 and Rating2 as well as between Rating1 and Rating3 was significant (both p<0.001), but the decrease between Rating2 and Rating3 was not (p=0.348).

**Figure 5:**
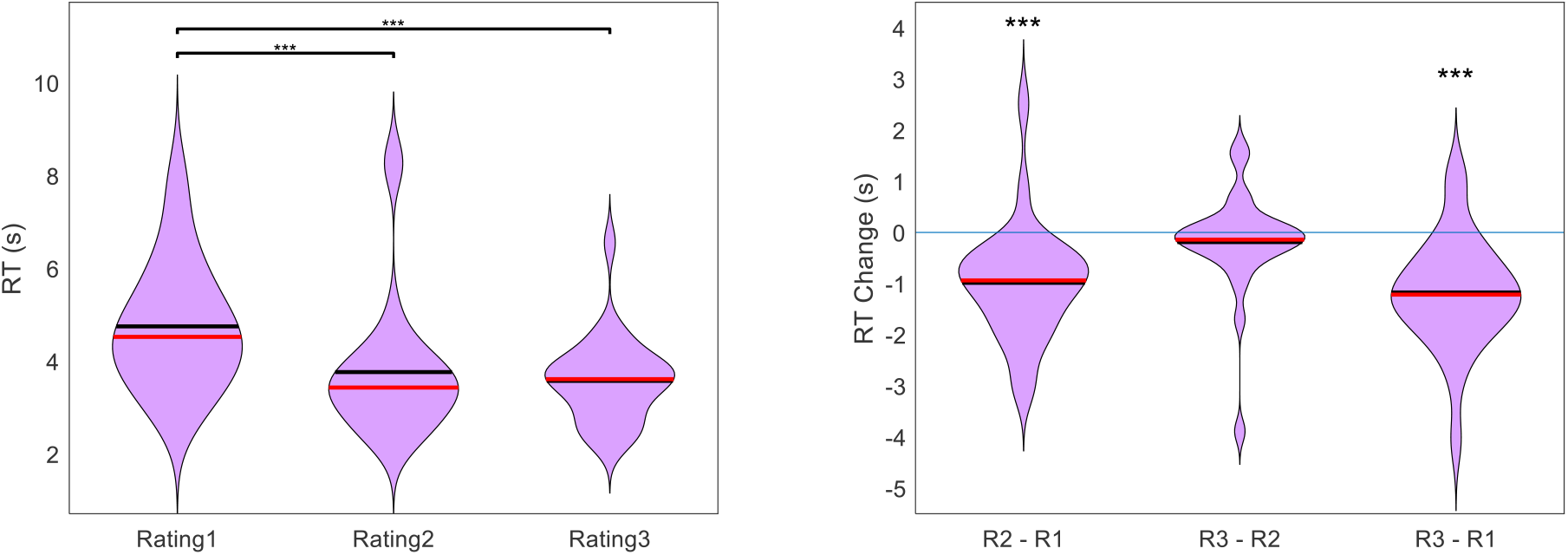
Across participants, RT (across items) decreased with each additional rating. Figure shows the cross-participant mean of within-participant mean RT across trials, separate for each rating task. (black lines indicate means, red lines indicate medians, significance stars indicate: * p<0.05, ** p<0.01, *** p<0.001)

We wondered if the decrease in RT over the course of the experiment (i.e., from Rating1 to Rating2 to Rating3) could simply be due to a habituation to the task. If this were true, RT would not only decrease across sections, but also within sections. We thus tested for a correlation between RT and trial number, within each rating section for each participant. Across participants, we found a mean correlation of −0.183 for Rating1 (p<0.001) and −0.079 for Rating2 (p=0.001), but no reliable correlation for Rating3 (mean = −0.033, p=0.238). In order to determine whether the decrease in RT from Rating1 to Rating2 (reported above) was actually to due certainty gains and not merely habituation, we split Rating1 into first half and second half trials. We then tested for a change in RT from Rating1 to Rating2 separately for first and second half trials, for each participant. There remained a significant decrease for both halves (first half mean RT change = - 1.20s, p<0.001; second half mean RT change = −0.78s, p<0.001).

#### Hypothesis 3: Certainty should negatively correlate with response time

For Rating1, the across-participant mean correlation between certainty and RT was negative, as predicted (mean Spearman’s rho = −0.104, p=0.004). For Rating2, there was no statistically significant correlation between certainty and RT (mean Spearman’s rho = 0.016, p=0.539). For Rating3, there was actually a positive correlation between certainty and RT (mean Spearman’s rho = 0.082, p=0.009).

Recalling that overall RT decreased across rating sections, we thought that this might have hidden the inherent relationship between certainty and RT. The idea is as follows. Initially (i.e., for Rating1), some items are evaluated with high certainty, others with low certainty. The high certainty evaluations are reported faster than the low certainty evaluations, thereby establishing the negative correlation between certainty and RT. Eventually (i.e., for Rating2 and Rating3), low-certainty evaluations become more certain (and thus more quickly evaluated). But, high-certainty evaluations remain certain, and there is not much room for an increase in certainty for these evaluations. Therefore, when averaging across the entire set of items, this would cause an overall increase in certainty as well as an overall decrease in RT. This could deteriorate the initial relationship between certainty and RT, as the set of items in effect shifts towards similarity (i.e., high certainty and low RT). To test this idea, we examined the evolution of certainty and RT on an item-by-item basis. We first calculated, for each participant and each item, the change in both the certainty and the RT from Rating1 to Rating2, and then calculated the within-participant correlation between those variables across items. Across participants, the correlation between Certainty Change and RT Change from Rating1 to Rating2 was indeed negative (mean Spearman’s rho = −0.051, p=0.031). We then repeated this same analysis using the differences from Rating2 to Rating3, and from Rating1 to Rating3. These correlations were both negative as well, although the former was not significant (mean Spearman’s rho = −0.022, p=0.235; mean Spearman’s rho = −0.053, p=0.015; two-sided t-tests). (see Figure 6).

**Figure 6:**
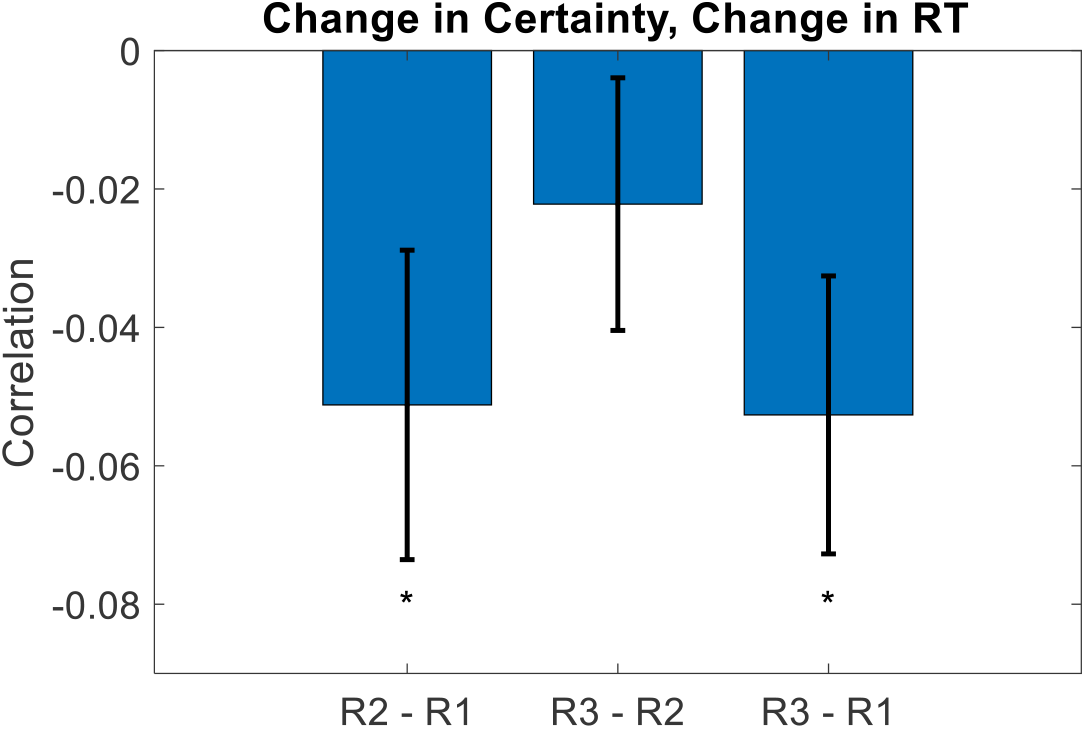
Across participants, the change in value rating certainty negatively correlated with the change in value rating RT. The idea here is that on an item-by-item basis, as certainty increases (across repeated ratings), it takes the DM less time to decide upon a rating estimate. (error bars represent s.e.m., significance stars represent: * p<0.05, ** p<0.01, *** p<0.001)

### Study 2

With Study 1, we demonstrated the reliability of our experimental measures of certainty regarding subjective value estimates. With Study 2, we seek to further demonstrate the importance of such measures by establishing their instrumental role in the decision making process.

#### Hypothesis 1: Choices will be more stochastic when value certainty is lower

Value-based choice is primarily a function of the difference in the value estimates of the different options in the choice set. The farther apart the value estimates are, the more likely it is that the higher-rated item will be chosen; the closer together the value estimates are, the more likely it is that the choice will appear to be random. Indeed, our data follow this pattern. For each participant, we performed a logistic regression of choices against the difference in value ratings of the paired options (choice = beta0 + beta1*dV + ε). We found that this function fit the data well above chance level, with a cross-participant average balanced accuracy of 77% (p<0.001, two-sided t-test). Across participants, there was no inherent bias for one side over the other (mean beta0 −0.036, p=0.350) and there was a significant positive inverse temperature parameter (mean beta1 = 0.077, p<0.001).

What would be more interesting, however, would be to see how value estimate certainty influences this choice model. We thus performed a similar logistic regression, for each participant, except this time we also included an indicator variable that took the value of 1 if the value certainty of a particular choice pair was greater than the median for that participant, and 0 otherwise (choice = beta0 + beta1*dV + beta2*I*dV + ε). Balanced accuracy remained at 77% (p<0.001). As with the previous model, there was no bias (mean beta0 = −0.035, p=0.356) and the inverse temperature for value difference was positive and significant (mean beta1 = 0.077, p<0.001). Notably, the regression coefficient for the interaction of value difference and the high certainty indicator (i.e., the increase in choice precision between low and high value certainty trials) was positive but only marginally significant (mean beta2 = 0.042, p=0.108). (See Figure 7.)

**Figure 7:**
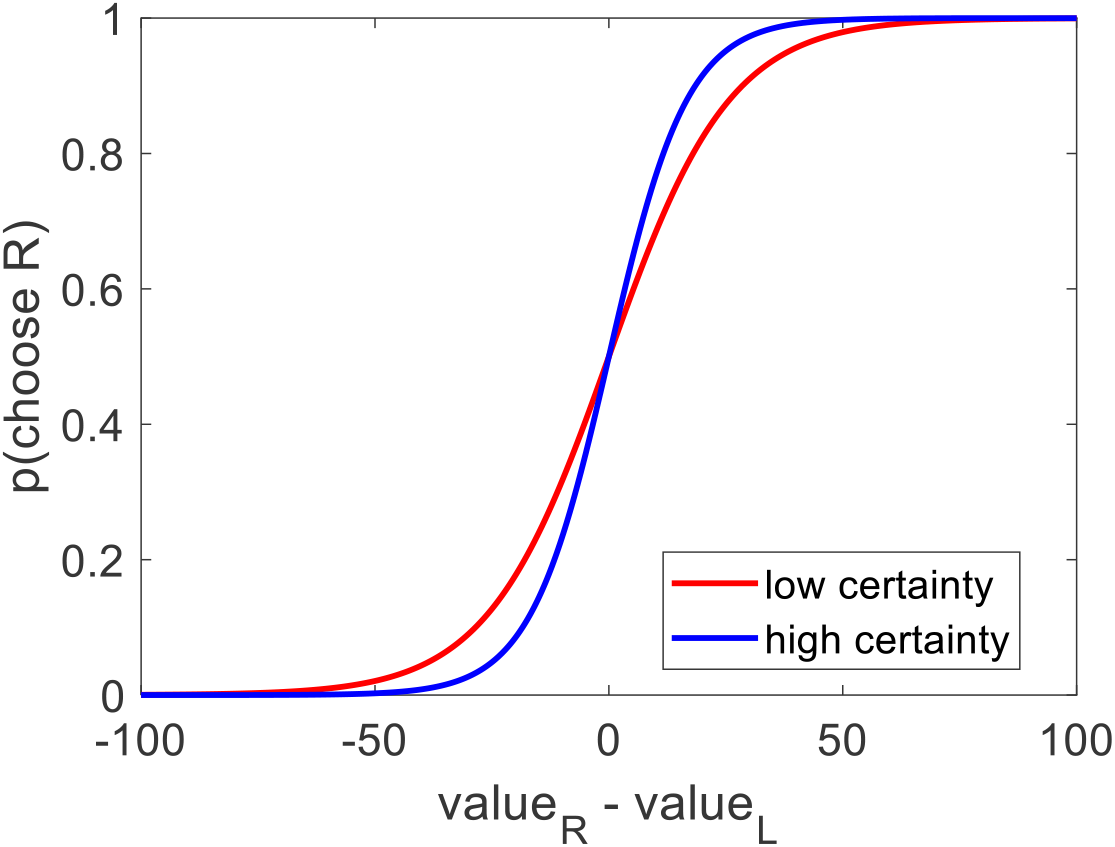
Across participants, the probability of choosing the option on the right increased as a function of the value estimate difference (right option – left option). In particular: choices that were made between options with low value certainty (red curve, within subject median split) were more stochastic than choices that were made between options with high value certainty (blue curve) (left plot).

#### Hypothesis 2: Options with higher value certainty will be chosen more often

We posited that choices might be partially determined by how certain the individual value estimates for each option were. We thus wondered how well choice could be predicted using the difference in value certainty alone, without considering the difference in value estimates themselves. For each participant, we ran a logistic regression of choices against the difference in value estimate certainty (choice = beta0 + beta1*dC + ε). Balanced accuracy was lower under this model, as expected, but it was still well above chance level (cross-participant mean = 59%, p<0.001). Again, there was no bias (mean beta0 = −0.044, p=0.134). The inverse temperature for value certainty difference was positive and significant, as expected (mean beta1 = 0.326, p<0.001) (see Figure 8). This shows that choices can indeed be predicted by the difference in the value certainty of the options under consideration, without directly examining the difference in the value estimates themselves.

**Figure 8:**
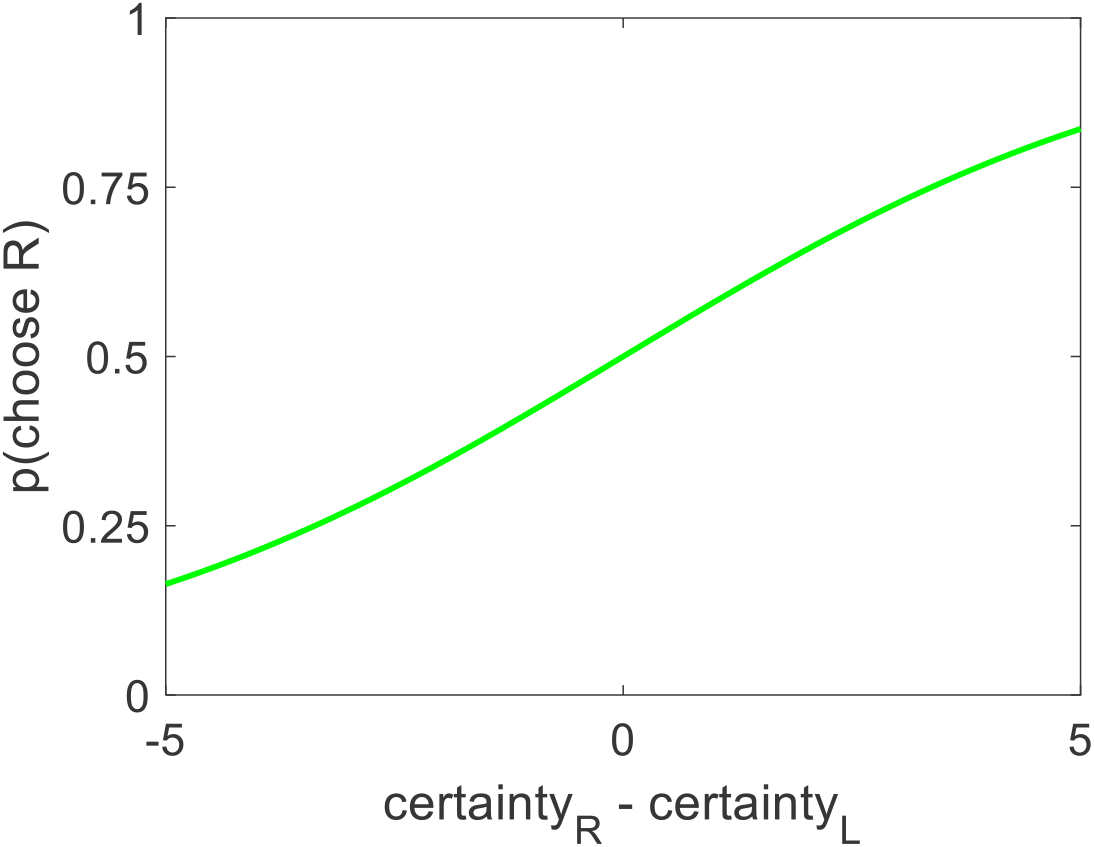
Across participants, the probability of choosing the option on the right increased as a function of the value certainty difference (right option – left option).

Although we showed that choice could be predicted by value certainty even without considering value estimate, we realized that there is generally a strong relationship between those two variables. Supporting this notion, we found that value certainty correlated positively with value estimate (mean Spearman’s rho = 0.254, p<0.001). Moreover, there was a clear u-shaped relationship between value estimate and value certainty (see Figure 9). We note, however, that the value certainty reports carried additional information beyond the value estimate ratings themselves. The data clearly show that whereas very high or very low value estimates almost always correspond to very high certainty, mid-range value estimates do not necessarily correspond to relatively low certainty. It seems that sometimes participants estimated an item’s value to be mid-range because they were not certain about its true value, but other times they were quite certain that its value was mid-range. This shows that value certainty is partially constrained, but not fully determined, by value estimate itself.

**Figure 9:**
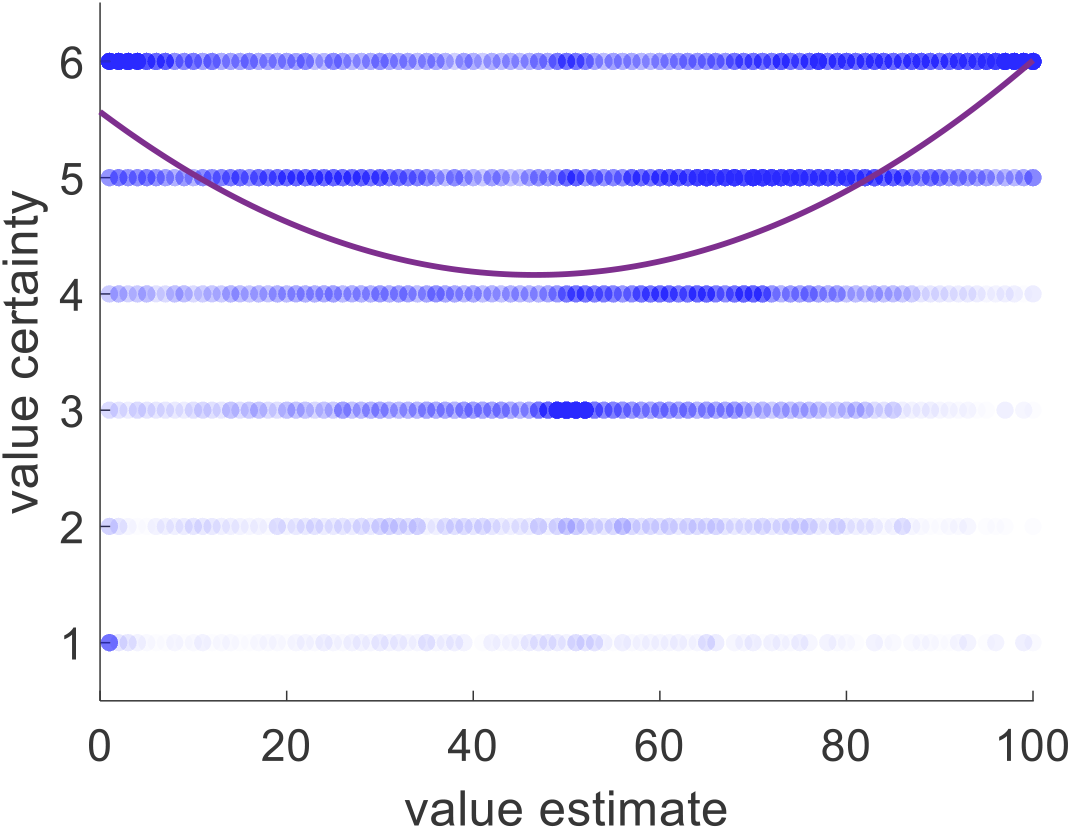
Value certainty is related to value estimate, with both a linear effect and a quadratic one. Blue dots represent one item for one participant for one rating session. Purple curve represents the best linear + quadratic fit across all trials and all participants.

Exploring further, we wondered if the predictive power of value certainty might be driven entirely by its relationship with value estimate. That is, we wanted to check if the information contained in the value certainty reports beyond what they convey about the value estimates themselves would be useful in predicting choice. We predicted that, all else equal on a particular trial, the option with the higher value estimate certainty would be the chosen option. To test this, we ran, for each participant, a logistic regression of choices against the difference in value estimate ratings as well as the difference in value certainty reports (choice = beta0 + beta1*dV + beta2*dC + ε). Prior to running the regression, we first z-scored value estimate ratings and value certainty reports separately for each participant. Balanced accuracy remained at 77% (p<0.001). As with the previous models, there was no bias (mean beta0 = −0.042, p=0.231), and the inverse temperature for value difference was positive and significant (mean beta1 = 1.626, p<0.001). Notably, the inverse temperature for certainty difference was also positive and significant (mean beta2 = 0.185, p<0.001). (See Figure 10.) The regression function we used orthogonalizes regressors sequentially when calculating beta weights (i.e., here dC was orthogonalized to dV), so the impact of dC was truly separate from the impact of dV. This suggests that not only did the participants consider the difference in value estimates when choosing their preferred options, but they also considered the difference in value certainty irrespective of the value estimates.

**Figure 10:**
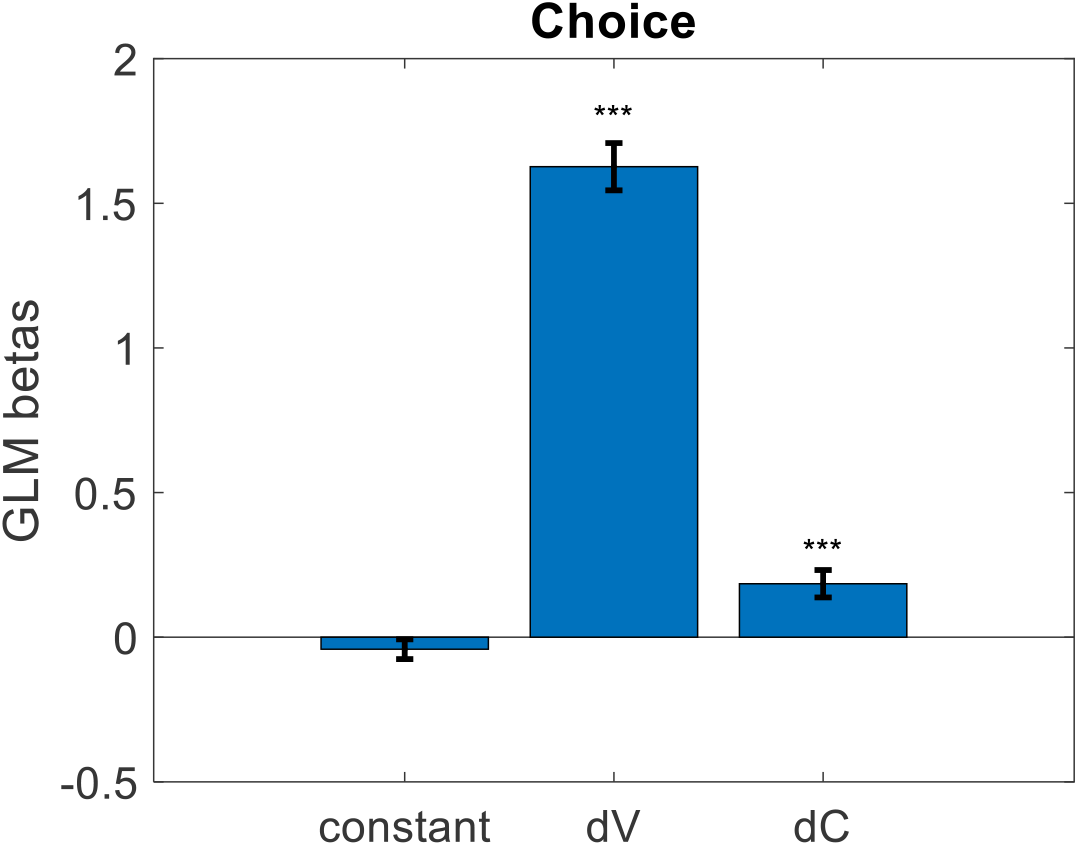
Cross-participant mean beta weights from GLM logistic regression of trial-by-trial value estimate difference (dV) and value certainty difference (dC) onto choice. (error bars represent s.e.m.)

#### Replication Results

After testing our hypotheses, we next performed a series of analyses to try to replicate previously reported results showing how value estimate certainty impacts a variety of dependent variables during choice (Lee and Daunizeau, 2020a, 2020b). Specifically, we checked whether choice confidence, response time, or choice-induced preference change changed as a function of value estimate certainty. For our measure of choice-induced preference change, we used the spreading of alternatives, defined as the post-minus pre-choice rating for the chosen option minus the post-minus pre-choice rating for the rejected option. For each of the above dependent variables, we ran a linear regression using absolute value estimate difference (vDiff) and summed value estimate certainty (cSum) as regressors. We removed an average of five trials per participant for having outlier RT (RT > median + 3*MAD), and z-scored all variables within participant. For choice confidence, we found that both independent variables had positive beta weights, as predicted (mean for vDiff = 0.276, p<0.001; mean for cSum = 0.068, p=0.016; two-sided t-tests). For response time, we found that both independent variables had negative beta weights, as predicted, although only vDiff was significant (mean beta for vDist = −0.206, p<0.001; mean beta for cSum = −0.011, p=0.586; two-sided t-tests). For spreading of alternatives, we found that both independent variables had negative beta weights, as predicted (mean beta for vDiff: −0.277, p<0.001; mean beta for cSum: −0.045, p=0.048; two-sided t-tests). (See Figure 11.)

**Figure 11:**
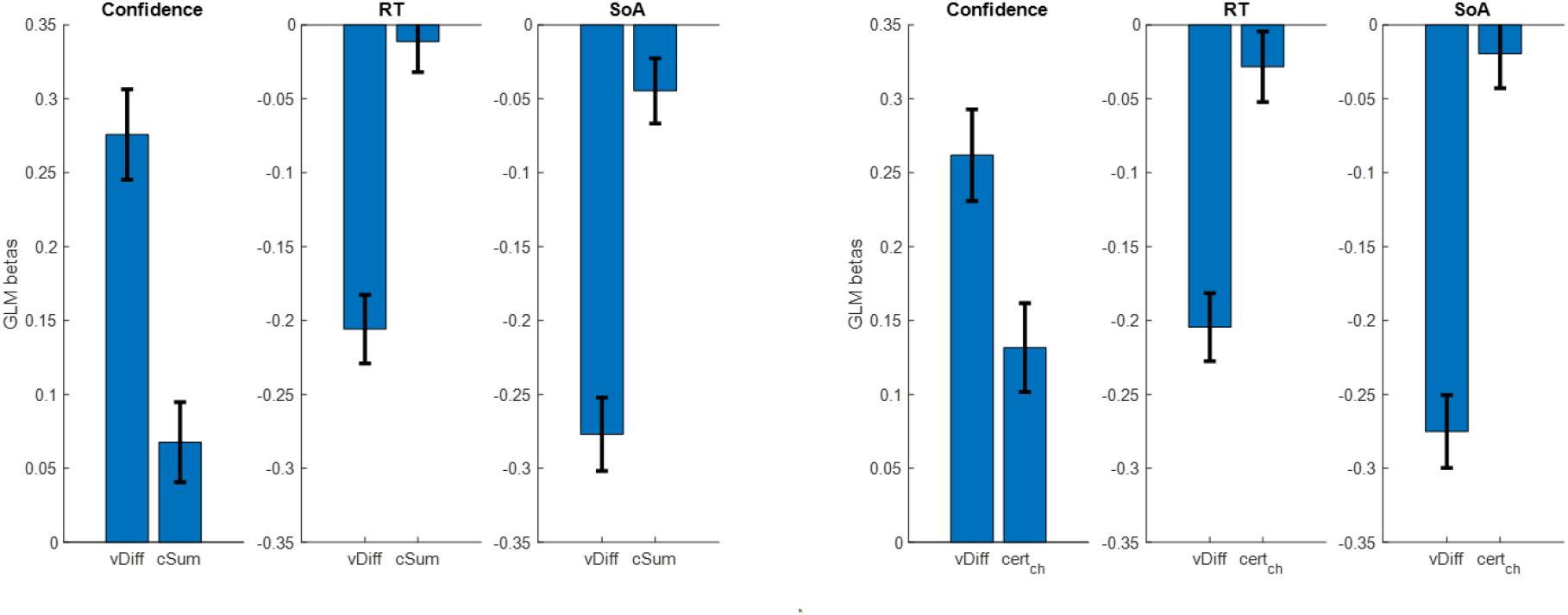
Cross-participant mean beta weights from GLM regressions of trial-by-trial absolute value estimate difference (vDiff) and summed value estimate certainty (cSum) onto choice confidence, choice response time, and spreading of alternatives (left set of plots); same, using certainty of only the chosen option (right set of plots). (error bars represent s.e.m.)

The use of cSum to represent the relevant aspect of value estimate certainty during choice deliberation was somewhat arbitrary. We therefore decided to examine other measures in the place of cSum, specifically: certainty of the chosen option (cert_ch_), certainty of the rejected option (cert_rej_), difference in certainty between the chosen and rejected options (cert_ch-rej_). For each participant, we repeated the same GLM regression as described above, replacing cSum with cert_ch_, cert_rej_, and cert_ch-rej_ in turn. We started with cert_ch_. For choice confidence, we found that both independent variables had positive beta weights, as predicted (mean for vDiff = 0.262, p<0.001; mean for cert_ch_ = 0.132, p<0.001; two-sided t-tests). For response time, we found that both independent variables had negative beta weights, as predicted, although only vDist was significant (mean beta for vDist = −0.205, p<0.001; mean beta for cert_ch_ = −0.028, p=0.240; two-sided t-tests). For spreading of alternatives, we found that both independent variables had negative beta weights, as predicted, although only vDiff was significant (mean beta for vDiff: −0.275, p<0.001; mean beta for cert_ch_: −0.020, p=0.408; two-sided t-tests). (See Figure 10 above.) The regression analyses using cert_rej_ did not yield significant beta weights for the certainty term (confidence: mean beta for vDiff: 0.275, p<0.001; mean beta for cert_rej_: −0.032, p=0.180; RT: mean beta for vDiff: −0.205, p<0.001; mean beta for cert_rej_: 0.010, p=0.659; SoA: mean beta for vDiff: −0.281, p<0.001; mean beta for cert_rej_: −0.039, p=0.108; two-sided t-tests). The regression analyses using cert_ch-rej_ yielded similar results as when using cert_ch_, though with slightly lower beta weights and slightly larger p-values (confidence: mean beta for vDiff: 0.263, p<0.001; mean beta for cert_ch-rej_: 0.116, p<0.001; RT: mean beta for vDiff: −0.204, p<0.001; mean beta for cert_ch- rej_: −0.021, p=0.402; SoA: mean beta for vDiff: −0.282, p<0.001; mean beta for cert_ch-rej_: 0.021, p=0.460; two-sided t-tests).

## DISCUSSION

In this study, we have demonstrated the reliability of multiple measures of subjective value estimate certainty, including self-reports, rating consistency, and response time. We have also demonstrated the important role that value estimate certainty plays in choice itself, including its positive impact on choice consistency and choice confidence, as well as its negative impact on response time and choice-induced preference change. We might suggest that any contemporary or future model of value-based decision making (and arguably, all types of decision making) should consider including some measure of value estimate certainty for each of the options in the choice set. At the present time, the only choice model that we are aware of that explicitly includes a variable to represent value estimate certainty is the Metacognitive Control of Decisions (MCD) presented by Lee and Daunizeau (2020a). This feature alone sets the MCD model apart from the plethora of alternative models that abound in the literature. Yet it would not be reasonable to claim that one class of model is inherently better than another simply because the alternative failed to consider an important variable. Rather, we propose that the popular models that already exist in the literature should be expanded to include value rating certainty. Only then can a more complete and fair model comparison be made, and only then will we be able to reach a better understanding of the cognitive mechanisms of decision making.

In particular, we call upon proponents of the so-called accumulation-to-bound models, such as the Race Model (RM) and the Drift-Diffusion Model (DDM), to strongly consider revising their models to include value estimate certainty. As it stands, most such models completely exclude the possibility of item-specific certainty. These models typically (or exclusively) account for stochasticity in the choice deliberation process at the system level, rather than at the option level. This means that such models can explain or predict variations in observed behavior that are dependent on choice context (e.g., clarity of perception, mental workload), but not on the composition of the choice set itself. Given that stochasticity is one of the fundamental components of evidence accumulation models (i.e., the diffusion parameter), it begs the question as to why the nature of the stochasticity has not been more thoroughly explored. A related line of work has indeed explored this question, concluding that uncertainty could spawn from noise in sensory processing, stochasticity in response selection, or imperfections in probabilistic inference (Drugowitsch et al, 2016). However, they did not discuss the possibility that choice options might have different degrees of certainty intentionally represented in the brain.

Recent work has proven that an accumulation-to-bound process such as that represented by the RM or DDM is an optimal policy, at least when optimality is defined as the maximization of reward in a series of sequential decisions with a limited amount of time (Tajima et al, 2016, 2019). These authors do indeed acknowledge the importance of certainty in their work, although it is not quite of the same nature as that which we described in our study. In the work of Tajima et al (2016, 2019), pre-choice certainty about an option refers to the prior belief that a DM has about the value distributions from which each option originates, rather than a belief about the value estimates of the options themselves. However, we have shown that item-specific pre-choice certainty is an important input to the choice deliberation process. Without a measure of item-specific certainty, such a model cannot account for variations in choice behavior when the different options originate from the same categorical set (e.g., snack foods). Tajima et al (2016) suggest that evidence accumulation serves to increase the certainty about the option values, but that the momentary evidence itself is uncertain. According to the authors, noise in the momentary evidence itself could originate both externally (e.g., the stochastic nature of stimuli, perceptual noise, ambiguity, incomplete knowledge) or internally (e.g., uncertain memory, value inference that extends over time) (Tajima et al, 2016). Here, the authors seem to pave the way for certainty measures that vary on an option-by-option basis, although they do not make this explicit in their work.

Other recent work has suggested that the evidence accumulation process illustrated by a DDM is influenced by attention (Sepulveda et al, 2020; Krajbich and Rangel, 2011; Krajbich et al, 2010). Specifically, it has been proposed that during choice deliberation, evidence accumulates at a higher rate for the option that is currently being gazed at, relative to the other option(s). This evidence might support value estimation directly (Krajbich and Rangel, 2011; Krajbich et al, 2010) or a more general goal-relevant information estimation (Sepulveda et al, 2020). However, neither of these models include value estimate certainty. Indeed, these models explicitly assume that both the prior uncertainty (i.e., variability in the environment from which the options originate) and the evidence uncertainty (i.e., stochasticity in samples drawn from probability distributions with fixed means) are identical across options. We have shown that such an assumption is not reasonable, and thus likely impedes model performance.

Furthermore, these authors do not specifically examine the role of value certainty in gaze allocation. If gaze fixation is what focuses information processing on each option in turn, it could be that a DM will be inclined to gaze more at options whose values are less certain. Similar to the concept of exploration in the classic exploration/exploitation dilemma (e.g., Daw et al, 2006), this gaze bias could be instrumental for the DM to make more informed choices (Callaway et al, 2020). Or, it could be that a DM will be inclined to gaze more at options whose values are *more* certain. Value certainty could be a measure of how reliable the information about the option is, which could bias gaze towards options with higher certainty. Further studies will be required to demonstrate the direction of the influence of value certainty on gaze patterns, but it is likely that it plays some important role in the decision process. Regardless of the direction, we hold that gaze duration should correlate with an increase in value estimate certainty.

## ACKNOWLEDGEMENTS

This manuscript has been released as a pre-print at https://www.biorxiv.org/content/10.1101/2020.06.16.155234v1.

## Supplementary Material

### Data Quality Check

#### Study 1

Before testing our hypotheses, we performed a number of simple data quality checks. First, we assessed the test-retest reliability of value ratings. For each participant, we thus measured the correlation between first rating (Rating1) and second rating (Rating2), across items. We found that ratings were generally consistent (median Spearman’s rho = 0.817). Most participants showed a correlation of greater than 60%. We then measured, for each participant, the correlation between Rating1 and Rating3. We found that ratings were generally consistent (median Spearman’s rho 0.818).

Next, we performed a similar assessment of the test-retest reliability of certainty reports. Before examining the certainty data, we first converted the qualitative reports to numbers (“not at all” = 1, “slightly” = 2, “somewhat” = 3, “fairly” = 4, “very” = 5, “extremely” = 6). For each participant we then measured the correlation between certainty for Rating1 (Certainty1) and certainty for Rating2 (Certainty2), across items. We found that certainty reports were generally consistent (median Spearman’s rho = 0.352), although much less so than value ratings. We then measured, for each participant, the correlation between Certainty1 and Certainty3, across items. We found that certainty reports were generally consistent (median Spearman’s rho = 0.353), although much less so than value ratings.

Because rating certainty is a key variable for testing our hypotheses, we needed to be sure that participants responded meaningfully to the rating certainty question. An analysis of how certainty correlates with our other variables of interest would not be possible where there is insufficient variability in the certainty data. For this reason, we calculated a score for how much variance each participant had across certainty reports. The median certainty report variance was 0.935, 0.882, and 0.599 for Rating1, Rating2, and Rating3, respectively. We deemed that there were no participants who were obvious outliers based on this score.

For each of the test-retest reliability measures described above, we searched for population outliers. We defined an outlier for a specific measure as a participant whose score was more than three median average deviations (MAD) away from the population median. This technique yielded three outlier participants based on the Rating1-Rating2 test-retest reliability scores, and eight outlier participants based on the Rating1-Rating3 test-retest reliability scores. There was one outlier with respect to Certainty1-Certainty2, and six with respect to Certainty1-Certainty3. Seven of the outlier participants were caught by more than one filter, which left us with a set of 11 total outlier participants for Study 1. We excluded these participants from our reported analyses.

#### Study 2

Before testing our hypotheses, we performed a number of simple data quality checks. First, we assessed the test-retest reliability of value ratings. For each participant, we thus measured the pairwise linear correlation between first rating (Rating1) and second rating (Rating2), across items. We found that ratings were generally consistent (median Spearman’s rho = 0.803, p<0.001).

Next, we performed a similar assessment of the test-retest reliability of certainty reports. Before examining the certainty data, we first converted the qualitative reports to numbers (“not at all” = 1, “slightly” = 2, “somewhat” = 3, “fairly” = 4, “very” = 5, “extremely” = 6). For each participant we then measured the pairwise linear correlation between certainty for Rating1 (Certainty1) and certainty for Rating2 (Certainty2), across items. We found that certainty reports were generally consistent (median Spearman’s rho = 0.344, p<0.001), although much less so than value ratings.

Because rating certainty is a key variable for testing our hypotheses, we needed to be sure that participants responded meaningfully to the rating certainty question. An analysis of how certainty correlates with our other variables of interest would not be possible where there is insufficient variability in the certainty data. For this reason, we calculated a score for how much variance each participant had across certainty reports. The median certainty report variance was 0.971 and 0.818 for Rating1 and Rating2, respectively. We deemed that there were no participants who were obvious outliers based on this score.

Finally, we checked whether choices were consistent with pre-choice ratings. For each participant, we performed a logistic regression of choices against the difference in value ratings of the paired options. We found that the balanced prediction accuracy was beyond chance level (mean 77%), indicating participants were performing the choice task properly.

For each of the test-retest reliability measures described above, we searched for population outliers. We defined an outlier for a specific measure as a participant whose score was more than three median average deviations (MAD) away from the population median. This technique yielded three outlier participants based on the Rating1-Rating2 test-retest reliability scores. We excluded these participants from our reported analyses.

##### Effect of confidence on choice consistency

We then explored a step further, postulating that choice confidence should modulate choice consistency (often referred to as accuracy). The idea is that for high confidence choices, the DM would more consistently distinguish the items, relative to low confidence choices. We thus performed a similar logistic regression as we did in our main analysis (see Figure 6 in Results), for each participant, except this time the indicator represented high choice confidence (within-participant median split) instead of value certainty (choice = logistic[beta0 + beta1*dV + beta2*Ind*dV]). Under this model, balanced accuracy was also 77% (p<0.001). Again, there was no bias (mean beta0 = −0.028, p=0.466), and the inverse temperature parameter remained positive and significant (mean beta1 = 0.065, p<0.001). Notably, the regression coefficient for the interaction of value difference and the high confidence indicator (i.e., the increase in choice precision between low and high confidence trials) was positive and significant (mean beta2 = 0.088, p<0.001) (see Figure S1). We thus confirmed a common observation that choice confidence and choice accuracy are closely linked.

**Figure S1:**
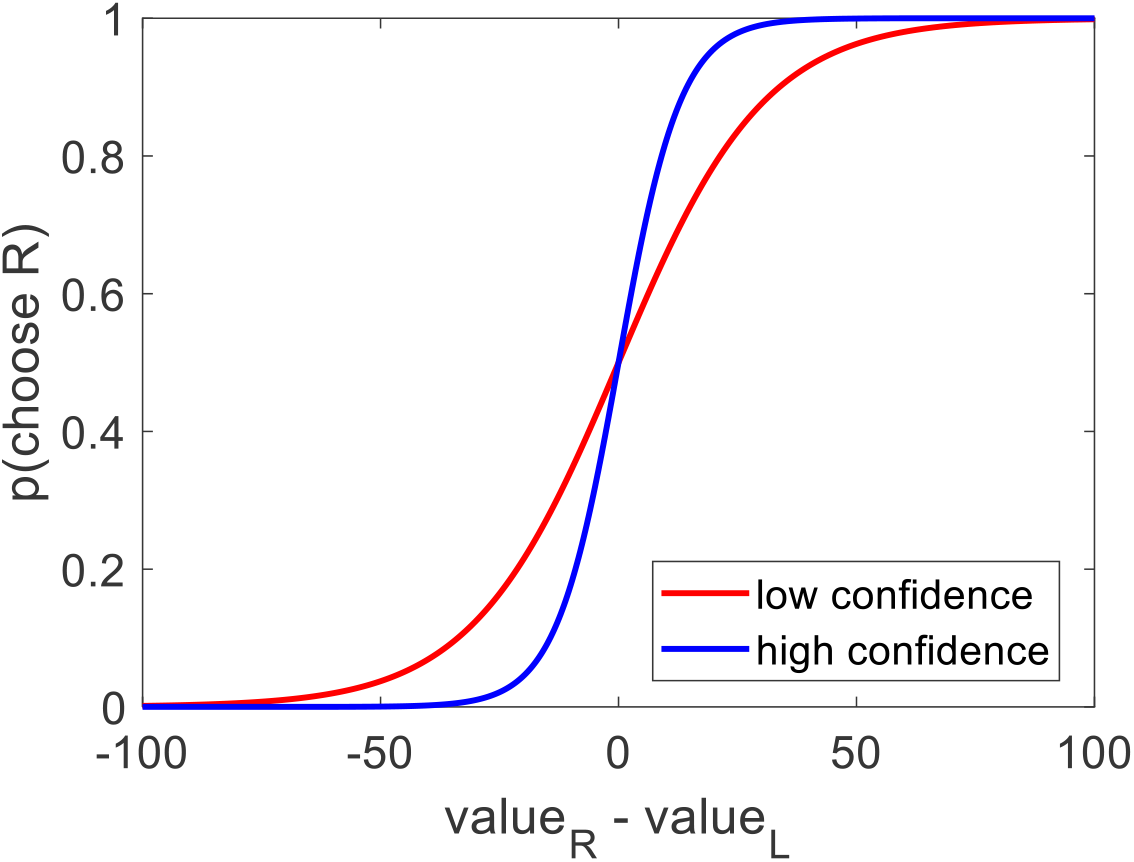
Across participants, the probability of choosing the option on the right increased as a function of the value estimate difference (right option – left option). In particular: choices that were made with low confidence (red curve, within subject median split) were more stochastic than choices that were made with high confidence (blue curve).

